# Simulation-based comparison of Biopharmaceutics Classification System and drug structure

**DOI:** 10.1101/631820

**Authors:** Tianyu Chen, Tianqiang Wu, Ningxi Li, Huanli Yin, Min Wu

**Affiliations:** Department of Pharmacy, Chengdu Medical College, No.783, Xindu Avenue, Xindu District, Chengdu, Sichuan Province, P.R.China

**Keywords:** Computer simulation, Molecular fingerprint, BCS, SVM

## Abstract

**Background:** The Biopharmaceutics Classification System (BCS), which classifies bioactive molecules based on solubility and permeability, is widely used to guide new drug development and drug formulation, as well as predict pharmacokinetics. Here we performed computer simulations to study correlations between a molecule’s structure and its BCS classification.

**Methods:** A total of 411 small molecules were assigned to BCS categories based on published drug data, and their Pybel-FP4 fingerprints were extrapolated. The information gain(IG) of each fingerprint was calculated and its characteristic structure analyzed. IG was calculated using multiple thresholds, and results were verified using support vector machine prediction, while taking into account the dose coefficient(0-0.1, 0.1-1, or>1). Structural functional features common to fingerprints of compounds in each type of BCS class were determined using computer simulations.

**Results:** BCS classes III and IV appear to share several structural and functional characteristics, including Secondary aliphaticamine, Michael_acceptor, Isothiourea, and Sulfonamide Sulfonic_derivatives.

**Conclusion:** We demonstrate that our approach can correlate characteristic fingerprints of small-molecule drugs with BCS classifications, which may help guide the development and optimization of new drugs.

## Introduction

The Biopharmaceutics Classification System (BCS) [1–2] was proposed in 1995 [3] to classify drugs into four categories based on their *in vitro* solubility and ability to be absorbed in the intestine (permeability):class I, high solubility/high permeability; class II, low solubility/high permeability; class III, high solubility/low permeability; and class IV, low solubility/low permeability [4] (Fig 1). The definition of high solubility according to the US Food and Drug Administration is that the highest dose of a single administration can be dissolved in 250 ml or less of an aqueous solution at 37℃ at pH1.0-7.5. Solubility and intestinal permeability have proven to be an adequate starting point for drug product development and regulation [5]. The role of BCS in drug development is facilitating biowaivers of *in vivo* bioequivalence studies [6]. At present, the BCS classification of drugs is achieved mainly experimentally, which requires a large amount of human, material, and financial resources, so a new method is urgently needed.

**Fig 1.**
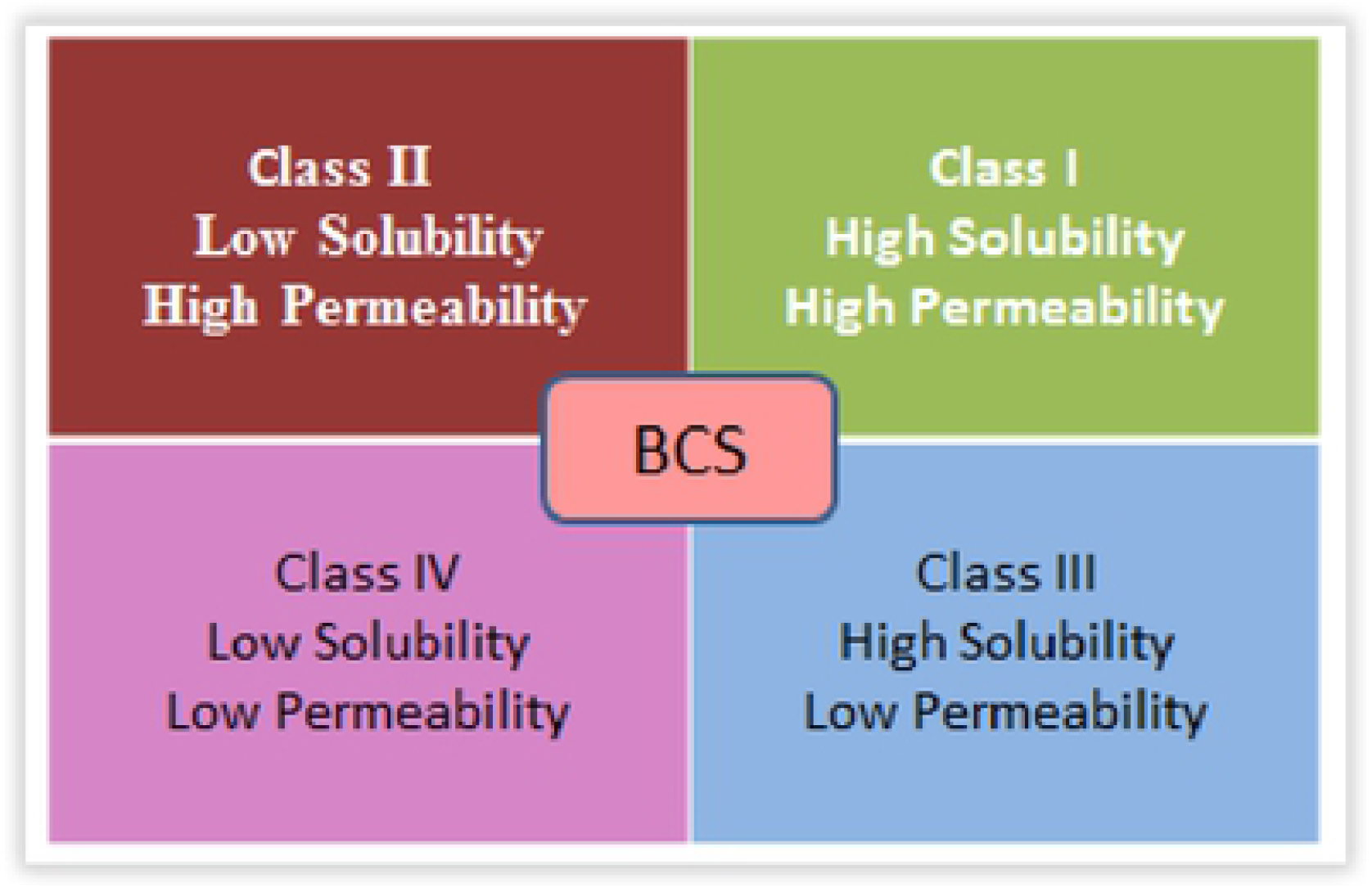
Biopharmaceutics Classification System. Schematic of the Biopharmaceutical classification system (BCS) detailing characteristics of each drug class.

Compared to experimental methods, computer-aided drug design(CADD) can reduce research and development costs and minimize the use of human and financial resources. Computer simulation is widely used in CADD: for example, FP4 and MACCS [7] molecular fingerprints [8] are used in new drug research. These fingerprints are developed to describe chemical structures in a chemical database. Meanwhile, computer simulation uses some evaluation indicators, such as information gain (IG), which refers to the weight of a fingerprint in this category[9], and frequency of a substructure (f) [10]. Even though a few studies of the application of computer simulation have been published recently, such as estimation of ADME (Absorption, Distribution, Metabolism, Excretion) properties and prediction of drug-induced liver toxicity [9–10], research on prediction of drug BCS classification based on structure has not been reported.

In this study, we combined BCS classification with computer simulation for the first time to identify the characteristic structures of each type of small-molecule drug. This approach may be useful for determining the BCS classification of new drugs. We used FP4 molecular fingerprints to describe the chemical structures of small-molecule drugs, and then calculated IG and f values to evaluate computer simulations. Furthermore, we used a Support-Vector Machine (SVM) [11–13] as the evaluation criteria for BCS classification accuracy (Fig 2). SVM is a two-class classification model for linear separable cases. For linear indivisible cases, samples with low-dimensional input space are transformed into high-dimensional feature spaces by nonlinear mapping algorithms so that samples can be classified linearly [14].

**Fig 2.**
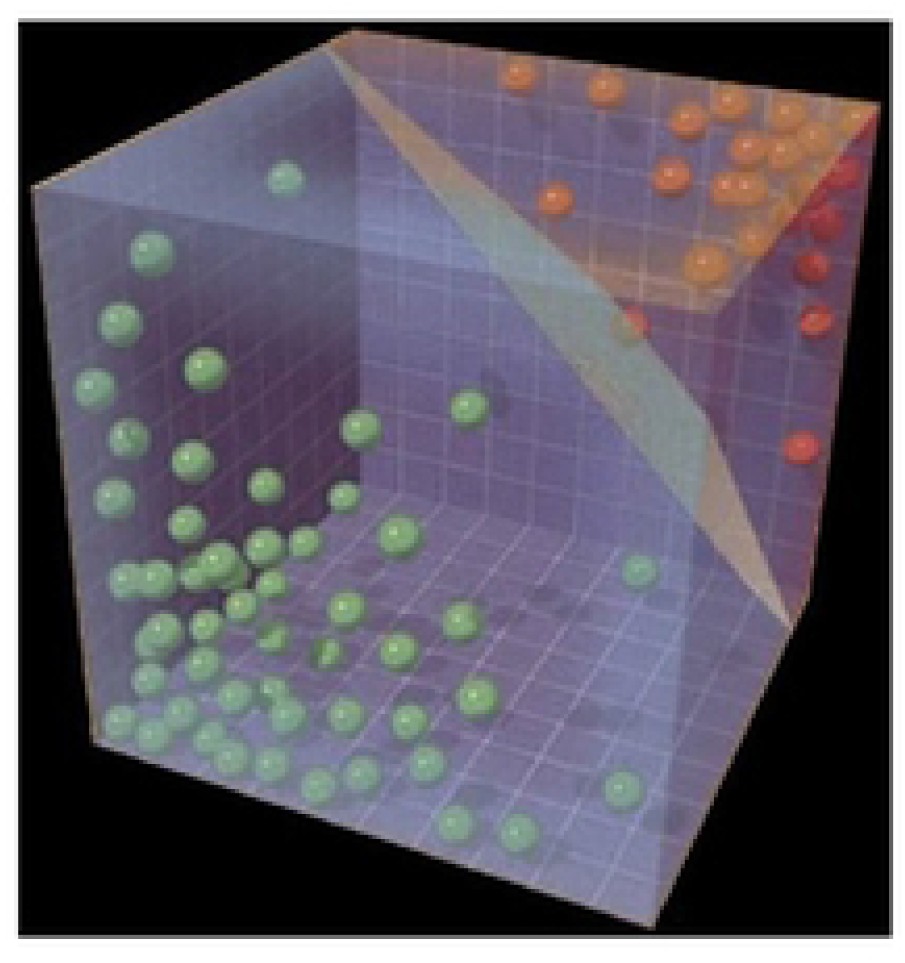
Support-Vector Machine schematic diagram. Schematic showing typical Support Vector Machine (SVM) data output. Red and green balls represent small-molecule drugs. The degree of difference, called a hyperplane, in the molecular fingerprints is given by the distance separatingthe red data points from the green point. The red or green data points closest to the hyperplane form thesupport vector.

In addition, we introduced the concept of dose coefficient F, which we define as the ratio of the molecular mass of each small-molecule drug to the mass in the maximum dose. Drug dissolution and absorption as well as the requirement for excipients are all critical factors in design of drugs in all BCS classes [15–17]. Dose size affects the absorption of the drug, and the maximum dose of the drug affects the solubility of the drug. Moreover, small-molecule drugs have different molecular masses. Therefore using the dose coefficient F may eliminate the influence of the maximum dose.

In these ways, the present study may help promote the development of new drugs and further development of existing drugs, and it may shorten the time-to-market for drugs.

## Materials and methods

### Establishment of database and molecular fingerprint analysis

A total of 359 small drug molecules were identified by the Provisional BCS Classification system (http://www.ddfint.net/search.cfm) (S1 Table). An additional 52 small drug molecules were identified in the literature using keyword search terms [18]. The molecular structures were downloaded (http://pubchem.ncbi.nlm.nih.gov/) for computer simulation. The BCS classification in the present study was the one recommended by the World Health Organization with CLogP [19] as the classification standard.

Molecular fingerprints were calculated by inserting the molecular structure into Open Babel software. The resulting .sdf files were entered into ChemDes (http://www.scbdd.com/fingerprints/index/), which calculated the Pybel-FP4 fingerprints[20]. The following formulas were used to calculate information entropy and IG value [9] for each molecular fingerprint:

Information entropy of all molecules in the entire database

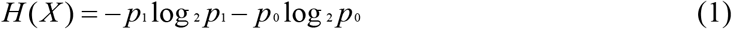

where p_1_ stands for the possibility of molecules in the first category, and p_0_ indicates the possibility of molecules in the second category.

The effect of a molecular fingerprint on the overall system

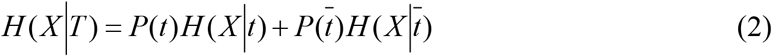

where P(t) stands for the probability that a molecular fingerprint will appear in the entire system, and 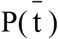 indicates the probability that a molecular fingerprint will not appear in the entire system.

Information entropy of a molecular fingerprint under high solubility conditions

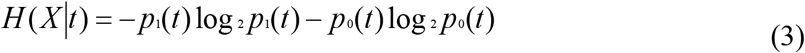

Information entropy of a molecular fingerprint under low solubility conditions

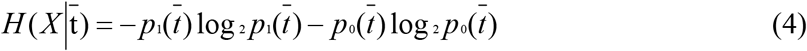

The influence of a molecular fingerprint on the entire molecule

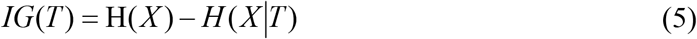

The larger the IG value, the greater is the effect of the structural composition on the entire molecular structure.

In this study, values of 1 equate to high solubility and 0 to low solubility.

Next, the frequency of a substructure (f) value was calculated [10]:

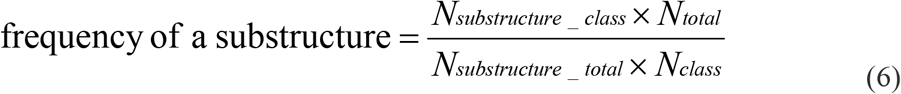

Both f and IG values were ordered from largest to smallest, and the first 20 values were selected and the common parts were taken as the characteristic molecular fingerprint to be determined. If the IG value of the top 20 molecular fingerprints was greater than 0.01, it indicated that the molecular fingerprint had a significant influence on the whole molecule, but a value below 0.01 meant the influence was small and the BCS category could not be clearly distinguished.

### SVM verification of BCS classification

To determine the accuracy of the SVM macros in differentiating small-molecule drugs into separate BCS classes, the classifications provided by the Provisional BCS website were used as a reference. IG values were extrapolated from molecular fingerprints and converted to binary file format. The binary FP4 molecular fingerprints were run through the SVM macros. Thresholds of 0, 0.001, 0.005, 0.01, or 0.02 were chosen based on solubility and permeability. The data were divided into training and test sets in the ratio of 1:4 based on the drug class assigned to each small molecule by the following macros: SP1(I-II), SP0(III-IV), PS1(I-III), PS0(II-IV), P(I, II-III, IV), S(I, III-II, IV).

A validation set [21] was created by examining the SVM output of the 359 small-molecule drugs from the known BCS classification as recognized by the World Health Organization. Data were derived as above, but thresholds were removed from the SVM macros. The accuracy of the output was compared to the known class designation of these molecules.

Every combination comparing one set to another was tested in the SVM software to calculate the accuracy of the classification. Due to the similarity between the training and test sets, the classification by the SVM software was considered accurate if it showed a value between 70% and 90%. A value less than 70% meant that the BCS classification was not complete enough to distinguish between the two types of data, while more than 90% meant excessive SVM training, such that the machine’s ability to generalize was insufficient. After the SVM was performed, molecular fingerprints were entered into SMARTS_InteLigand from Open Babel (http://www.scbdd.com/pybel_desc/fps-fp4/) [22] and the small-molecule drug structure was recreated based on fingerprint characteristics.

### Classification based on dose coefficient F

When the SVM verification result was outside the ideal percentage range, the accuracy of the computer simulation prediction was improved using secondary classification according to dose coefficient F. We defined the concept of dose coefficient F for the first time, which considers the ratio of the molecular mass of each small-molecule drug to the mass in the maximum dose. This classification was divided into the following categories: 0 to 0.1, 0.1 to 1, and >1.

### Verification of results

The obtained BCS feature structures were compared with those of small drug molecules with clear BCS classification in the BCS database to verify whether the results were accurate (Fig 3).

**Fig 3.**
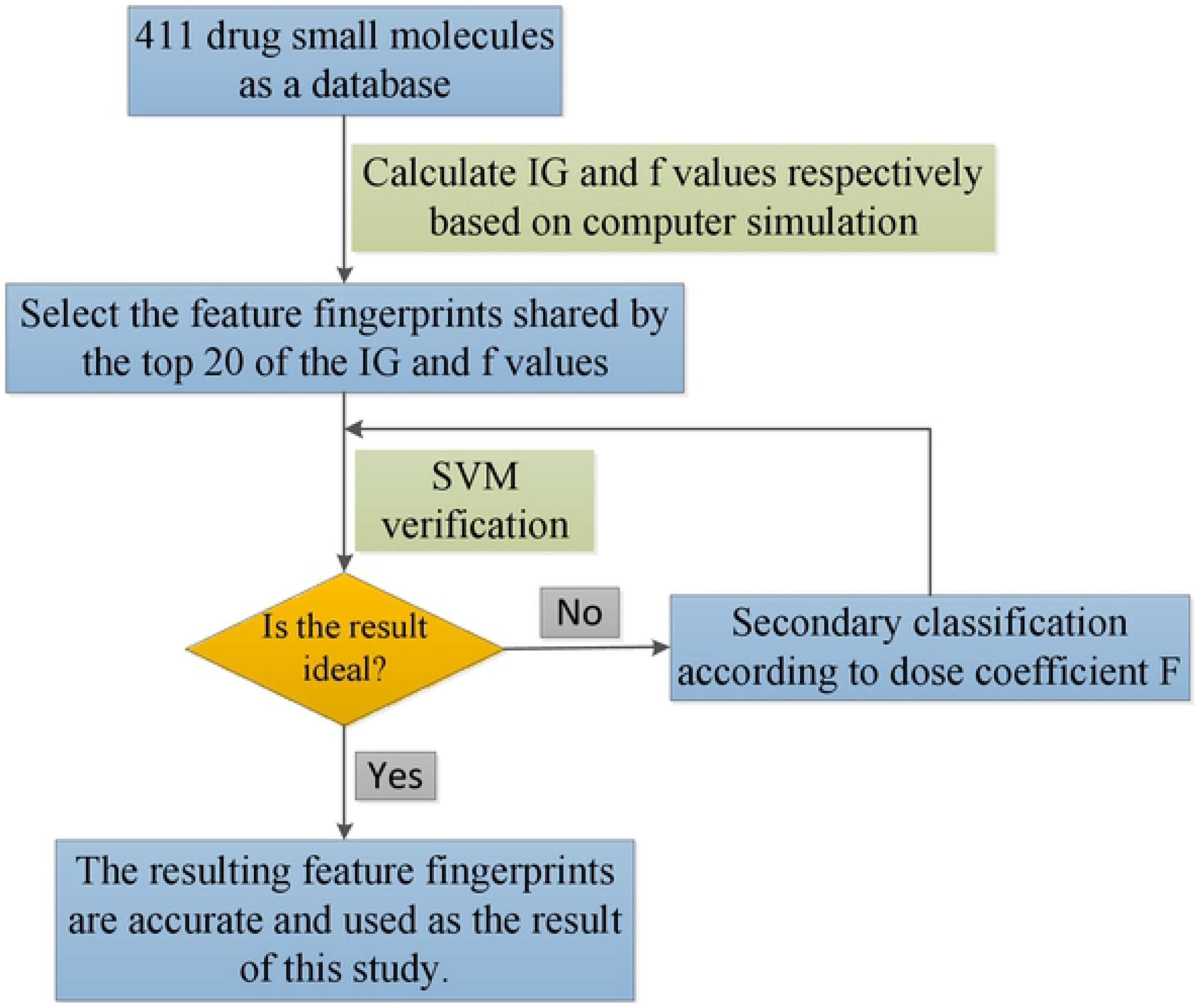
Experimental flowchart. Schematic describing the work flow of the study. Abbreviations: IG, Information gain; f, frequency of a substructure; SVM, Support Vector Machine.

## Results

### Molecular fingerprints sharing IG and f values

Preliminary screening was performed by identifying commonly shared molecular fingerprints from the lists containing the top 20 results with either the highest IG or f values (Tables 1-4 and S1 File). However, the common molecular fingerprints determined by IG and f values overlapped among BCS categories, making it impossible to assign molecules to a BCS category based solely on these values. Therefore, this study utilized SVM software to verify accuracy.

**Table 1.**
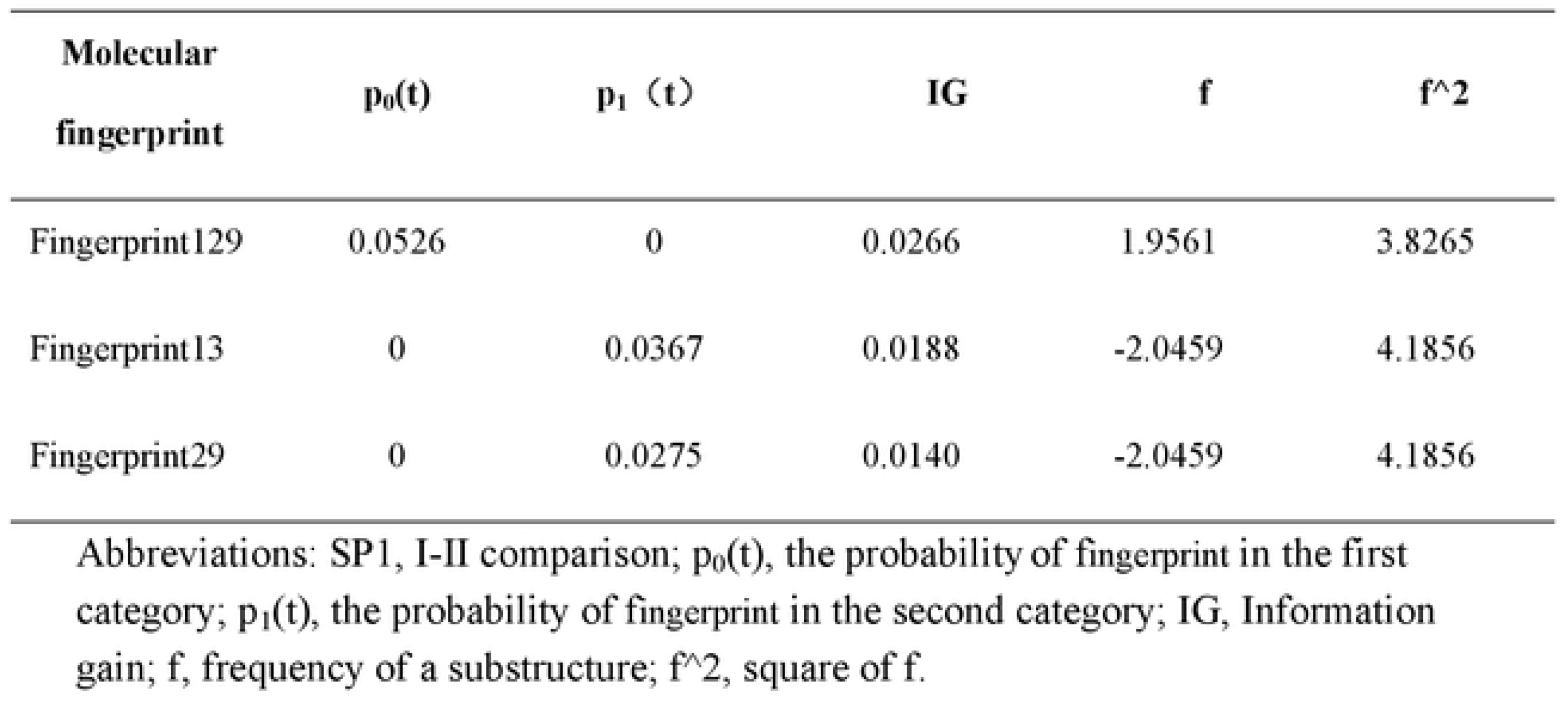
I-II comparison(SP1) result (S1 File. I-II Comparison(SP1)). Abbreviations: SP1, I-II comparison; p_0_(t), the probability of fingerprint in the first category; p_1_(t), the probability of fingerprint in the second category; IG, Information gain; f, frequency of a substructure; f^2, square of f.

**Table 2.** III-IV comparison(SP0) result (S1 File. III-IV Comparison(SP0)). Abbreviations: SP0, III-IV comparison; p_0_(t), the probability of fingerprint in the first category; p_1_(t), the probability of fingerprint in the second category; IG, Information gain; f, frequency of a substructure; f^2, square of f.

| **Molecular**<br>**fingerprint** | **$p_0(t)$** | **$p_1(t)$** | **IG** | **f** | **$f^2$** |
| --- | --- | --- | --- | --- | --- |
| Fingerprint211 | 0.0102 | 0.1667 | 0.0635 | -2.7381 | 7.4972 |
| Fingerprint214 | 0.0102 | 0.1667 | 0.0635 | -2.7381 | 7.4972 |
| Fingerprint183 | 0.0102 | 0.1429 | 0.0514 | -2.6531 | 7.0387 |
| Fingerprint55 | 0 | 0.0952 | 0.0511 | -3.3333 | 11.1111 |
| Fingerprint182 | 0.0102 | 0.1190 | 0.0398 | -2.5397 | 6.4500 |
| Fingerprint65 | 0.0408 | 0.1905 | 0.0390 | -1.7460 | 3.0486 |
| Fingerprint66 | 0.0408 | 0.1905 | 0.0390 | -1.7460 | 3.0486 |
| Fingerprint49 | 0.0102 | 0.0952 | 0.0287 | -2.3810 | 5.6689 |
| Fingerprint150 | 0.0102 | 0.0952 | 0.0287 | -2.3810 | 5.6689 |
| Fingerprint101 | 0.0714 | 0 | 0.0265 | 1.4286 | 2.0408 |
| Fingerprint200 | 0 | 0.0476 | 0.0252 | -3.3333 | 11.1111 |

**Table 3.** I-III comparison(PS1) result (S1 File. I-III Comparison(PS1)). Abbreviations: PS1, I-III comparison; p_0_(t), the probability of fingerprint in the first category; p_1_(t), the probability of fingerprint in the second category; IG, Information gain; f, frequency of a substructure; f^2, square of f.

| **Molecular**<br>**fingerprint** | **$p_0(t)$** | **$p_1(t)$** | **IG** | **f** | **$f^2$** |
| --- | --- | --- | --- | --- | --- |
| Fingerprint13 | 0 | 0.2857 | 0.1641 | -2.1633 | 4.6797 |
| Fingerprint41 | 0 | 0.0816 | 0.0433 | -2.1633 | 4.6797 |
| Fingerprint281 | 0 | 0.0816 | 0.0433 | -2.1633 | 4.6797 |
| Fingerprint229 | 0 | 0.0408 | 0.0213 | -2.1633 | 4.6797 |

**Table 4.** II-IV comparison (PS0) result (S1 File. II-IV Comparison(PS0)). Abbreviations: PS0, II-IV comparison; p_0_(t), the probability of fingerprint in the first category; p_1_(t), the probability of fingerprint in the second category; IG, Information gain; f, frequency of a substructure; f^2, square of f.

| **Molecular**<br>**fingerprint** | **$p_0(t)$** | **$p_1(t)$** | **IG** | **f** | **$f^2$** |
| --- | --- | --- | --- | --- | --- |
| Fingerprint17 | 0.0092 | 0.2143 | 0.0888 | -3.0972 | 9.5925 |
| Fingerprint28 | 0.0459 | 0.3095 | 0.0849 | -2.2117 | 4.8918 |
| Fingerprint65 | 0.0092 | 0.1905 | 0.0761 | -3.0418 | 9.2528 |
| Fingerprint66 | 0.0092 | 0.1905 | 0.0761 | -3.0418 | 9.2528 |
| Fingerprint128 | 0.0092 | 0.1190 | 0.0403 | -2.7651 | 7.6460 |
| Fingerprint24 | 0.0092 | 0.0952 | 0.0292 | -2.5991 | 6.7555 |
| Fingerprint55 | 0.0092 | 0.0952 | 0.0292 | -2.5991 | 6.7555 |
| Fingerprint37 | 0 | 0.0476 | 0.0248 | -3.5952 | 12.9257 |
| Fingerprint99 | 0 | 0.0476 | 0.0248 | -3.5952 | 12.9257 |
| Fingerprint129 | 0 | 0.0476 | 0.0248 | -3.5952 | 12.9257 |

### SVM prediction results

The accuracy of the BCS classification between the training and test sets was evaluated by SVM (Tables 5-10). When the IG threshold was 0.005 and only solubility or permeability was considered, the resulting SVM prediction was highly accurate (70-85%). In contrast, when drug classes were compared based on high or low solubility or high and low permeability, the accuracy of the SVM prediction was low, indicating large variability between the BCS populations. These data suggest that the current terms defining each drug class are too vague for this type of throughput and require additional information to improve classification accuracy.

**Table 5.** SVM predictive value under high permeability. Abbreviations: SP1, I-II comparison; SVM, Support-Vector Machine.

| **SP1(I-II)** | **Training (%)** | **Test (%)** |
| --- | --- | --- |
| 0 | 79.7753 | 71.1111 |
| 0.001 | 79.7753 | 62.2222 |
| 0.005 | 76.9663 | 73.3333 |
| 0.01 | 77.5281 | 68.8889 |
| 0.02 | 73.0337 | 73.3333 |

Abbreviations: SP1, I-II comparison; SVM, Support-Vector Machine.

**Table 6.** SVM predictive value under low permeability. Abbreviations: SP0, III-IV comparison; SVM, Support-Vector Machine.

| **SP0(III-IV)** | **Training (%)** | **Test (%)** |
| --- | --- | --- |
| 0 | 71.4286 | 64.2857 |
| 0.001 | 71.4286 | 64.2857 |
| 0.005 | 71.4286 | 71.4286 |
| 0.01 | 69.6429 | 71.4286 |
| 0.02 | 79.4643 | 67.8571 |

Abbreviations: SP0, III-IV comparison; SVM, Support-Vector Machine.

**Table 7.** SVM predictive value under high solubility. Abbreviations: PS1, I-III comparison; SVM, Support-Vector Machine.

| **PS1(I-III)** | **Training (%)** | **Test (%)** |
| --- | --- | --- |
| 0 | 78.2353 | 71.4286 |
| 0.001 | 76.4706 | 71.4286 |
| 0.005 | 80.0000 | 85.7143 |
| 0.01 | 85.2941 | 69.0476 |
| 0.02 | 80.5882 | 80.9524 |

Abbreviations: PS1, I-III comparison; SVM, Support-Vector Machine.

**Table 8.** SVM predictive value under low solubility. Abbreviations: PS0, II-IV comparison; SVM, Support-Vector Machine.

| **PS0(II-IV)** | **Training (%)** | **Test (%)** |
| --- | --- | --- |
| 0 | 71.9008 | 66.6667 |
| 0.001 | 72.7273 | 70.0000 |
| 0.005 | 73.5537 | 83.3333 |
| 0.01 | 76.8595 | 69.0476 |
| 0.02 | 75.2066 | 80.0000 |

Abbreviations: PS0, II-IV comparison; SVM, Support-Vector Machine.

**Table 9.** SVM predictive value compared with high permeability and low permeability. Abbreviations: P, I.II-III.IV comparison; SVM, Support-Vector Machine.

| **P(I.II-III.IV)** | **Training (%)** | **Test (%)** |
| --- | --- | --- |
| 0 | 75.8600 | 76.7100 |
| 0.001 | 61.3800 | 61.6400 |
| 0.005 | 61.0300 | 63.0100 |
| 0.01 | 63.4500 | 53.4200 |
| 0.02 | 61.7200 | 60.2700 |

Abbreviations: P, I.II-III.IV comparison; SVM, Support-Vector Machine.

**Table 10.**
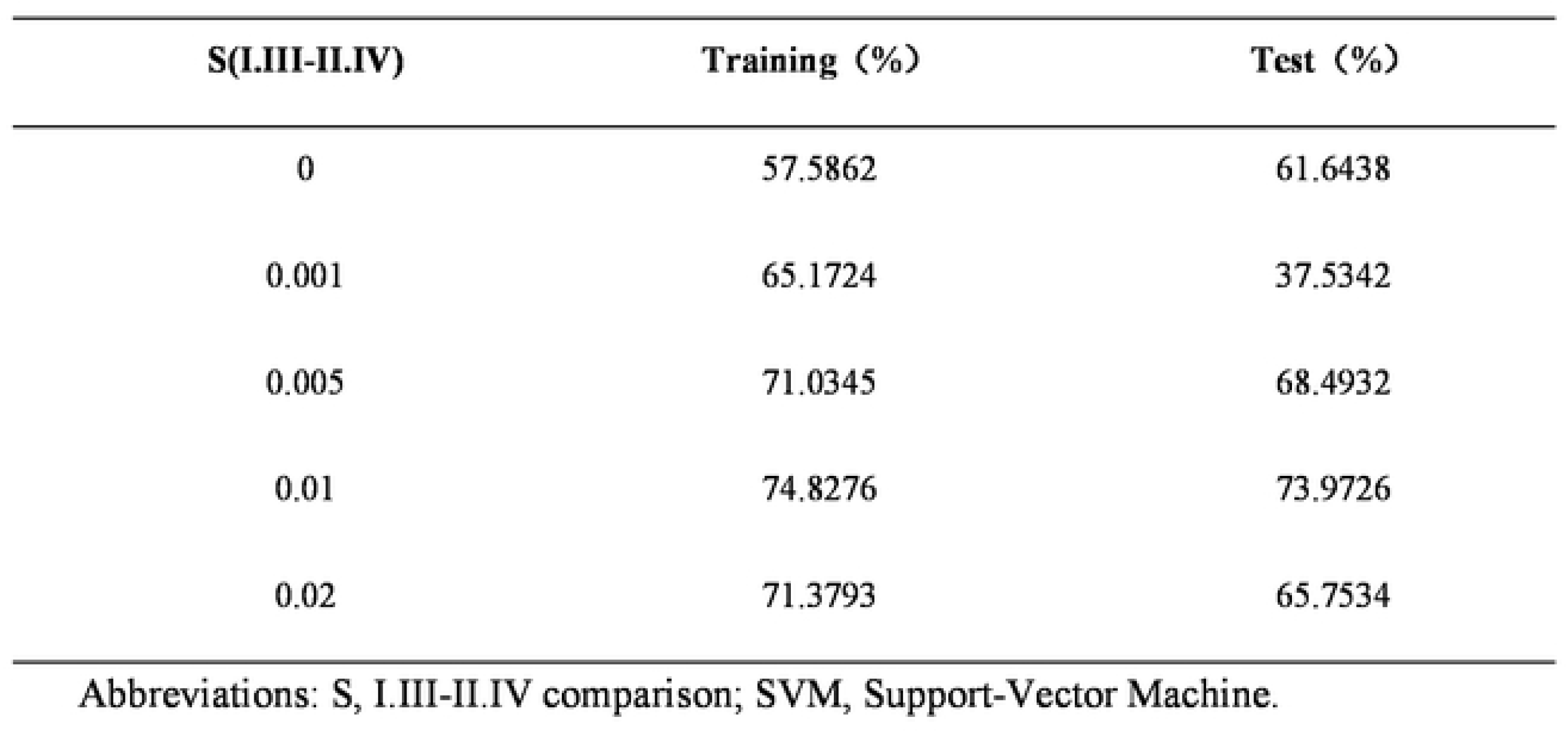
SVM predictive value compared with high solubility and low solubility. Abbreviations: S, I.III-II.IV comparison; SVM, Support-Vector Machine.

| **S(I.III-II.IV)** | **Training (%)** | **Test (%)** |
| --- | --- | --- |
| 0 | 57.5862 | 61.6438 |
| 0.001 | 65.1724 | 37.5342 |
| 0.005 | 71.0345 | 68.4932 |
| 0.01 | 74.8276 | 73.9726 |
| 0.02 | 71.3793 | 65.7534 |

Abbreviations: S, I.III-II.IV comparison; SVM, Support-Vector Machine.

### SVM prediction results based on dose coefficient F classification

The BCS classification was subdivided by the dose coefficient, and SVM prediction was performed (Tables 11-16 and S2 File). Compared to the SVM predictions without dose coefficient F, the SVM predictions with the coefficient of 0-0.1 improved BCS prediction accuracy when the SVM had a threshold of 0.01 or 0.02. Furthermore, when the dose coefficient was greater than 1, the accuracy of the SVM prediction improved within each threshold range. This indicated that the dose coefficient F affected the accuracy of the SVM prediction and the screening of the characteristic molecular fingerprint. Therefore, addition of the dose coefficient F can improve SVM accuracy.

**Table 11.** SVM value of I-II comparison after classification based on FP4 type BCS dose coefficient. Abbreviations: BCS, Biopharmaceutical classification system; SVM, Support-Vector Machine.

| Group | Threshold | Training (%) | Test (%) |
| --- | --- | --- | --- |
| Pybel.FP4-BCS-F-0-0.1-I-II | 0 | 70.0000 | 70.0000 |
|  | 0.001 | 70.0000 | 70.0000 |
|  | 0.005 | 71.2500 | 65.0000 |
|  | <u>0.01</u> | <u>68.7500</u> | <u>75.0000</u> |
|  | 0.02 | 72.5000 | 60.0000 |
| Pybel.FP4-BCS-F-0.1-1-I-II | 0 | 61.2500 | 78.4211 |
|  | 0.001 | 61.2500 | 52.6316 |
|  | 0.005 | 73.7500 | 78.9474 |
|  | 0.01 | 68.7500 | 73.6842 |
|  | <u>0.02</u> | <u>80.0000</u> | <u>89.4737</u> |
| Pybel.FP4-BCS-F-1-I-II | 0 | 82.3529 | 80.0000 |
|  | <u>0.001</u> | <u>82.3529</u> | <u>80.0000</u> |
|  | 0.005 | 76.4706 | 100.0000 |
|  | 0.01 | 76.4706 | 100.0000 |
|  | 0.02 | 88.2353 | 60.0000 |

**Table 12.** SVM value of I-III comparison after classification based on FP4 type BCS dose coefficient. Abbreviations: BCS, Biopharmaceutical classification system; SVM, Support-Vector Machine.

| Group | Threshold | Training (%) | Test (%) |
| --- | --- | --- | --- |
| Pybel.FP4-BCS-F-0-0.1-I-III | 0 | 61.7284 | 83.3333 |
|  | 0.001 | 66.6667 | 66.6667 |
|  | 0.005 | 69.1358 | 58.3333 |
|  | 0.01 | 64.1975 | 75.0000 |
|  | 0.02 | 67.9012 | 62.5000 |
| Pybel.FP4-BCS-F-0.1-1-I-III | 0 | 80.3030 | 62.5000 |
|  | <u>0.001</u> | <u>80.3030</u> | <u>81.2500</u> |
|  | 0.005 | 81.8182 | 56.2500 |
|  | 0.01 | 83.3333 | 68.7500 |
|  | 0.02 | 83.3333 | 93.7500 |
| Pybel.FP4-BCS-F-1-I-III | 0 | 85.7143 | 83.3333 |
|  | 0.001 | 90.4762 | 66.6667 |
|  | <u>0.005</u> | <u>85.7143</u> | <u>83.3333</u> |
|  | 0.01 | 90.4762 | 66.6667 |
|  | 0.02 | 80.9524 | 100.0000 |

**Table 13.** SVM value of II-IV comparison after classification based on FP4 type BCS dose coefficient. Abbreviations: BCS, Biopharmaceutical classification system; SVM, Support-Vector Machine.

| Group | Threshold | Training (%) | Test (%) |
| --- | --- | --- | --- |
| Pybel.FP4-BCS-F-0-0.1-II-IV | 0 | 83.3333 | 62.5000 |
|  | 0.001 | 83.3333 | 62.5000 |
|  | 0.005 | 76.6667 | 87.5000 |
|  | <u>0.01</u> | <u>76.6667</u> | <u>87.5000</u> |
|  | 0.02 | 80.0000 | 75.0000 |
| Pybel.FP4-BCS-F-0.1-1-II-IV | 0 | 71.8750 | 81.2500 |
|  | <u>0.001</u> | <u>73.4375</u> | <u>75.0000</u> |
|  | 0.005 | 43.4375 | 75.0000 |
|  | 0.01 | 73.4375 | 75.0000 |
|  | 0.02 | 75.0000 | 81.2500 |
| Pybel.FP4-BCS-F-1-II-IV | 0 | 59.2593 | 57.1429 |
|  | 0.001 | 70.3704 | 42.8571 |
|  | 0.005 | 70.3704 | 57.1429 |
|  | <u>0.01</u> | <u>81.4815</u> | <u>71.4826</u> |
|  | 0.02 | 70.3704 | 42.8571 |

**Table 14.** SVM value of III-IV comparison after classification based on FP4 type BCS dose coefficient. Abbreviations: BCS, Biopharmaceutical classification system; SVM, Support-Vector Machine.

| Group | Threshold | Training (%) | Test (%) |
| --- | --- | --- | --- |
| Pybel.FP4-BCS-F-0-0.1-III-IV | 0 | 83.8710 | 75.0000 |
|  | 0.001 | 87.0968 | 66.6667 |
|  | <u>0.005</u> | <u>77.4194</u> | <u>91.6667</u> |
|  | 0.01 | 90.3226 | 58.3333 |
|  | 0.02 | 100.0000 | 33.3330 |
| Pybel.FP4-BCS-F-0.1-1-III-IV | 0 | 66.0000 | 76.9231 |
|  | 0.001 | 68.0000 | 61.5385 |
|  | 0.005 | 74.0000 | 69.2308 |
|  | 0.01 | 74.0000 | 69.2308 |
|  | <u>0.02</u> | <u>72.0000</u> | <u>76.9231</u> |
| Pybel.FP4-BCS-F-1-III-IV | 0 | 58.0645 | 62.5000 |
|  | 0.001 | 61.2903 | 50.0000 |
|  | 0.005 | 61.2903 | 50.0000 |
|  | 0.01 | 74.1935 | 100.0000 |
|  | 0.02 | 64.5161 | 75.0000 |

**Table 15.** SVM values of I, II-III, IV comparison after classification based on FP4 type BCS dose coefficient. Abbreviations: BCS, Biopharmaceutical classification system; SVM, Support-Vector Machine.

| Group | Threshold | Training (%) | Test (%) |
| --- | --- | --- | --- |
| Pybel.FP4-BCS-F-0-0.1-I.II-III.IV | 0 | 67.5439 | 79.3103 |
|  | 0.001 | 69.2982 | 72.4138 |
|  | 0.005 | 72.8070 | 58.6207 |
|  | 0.01 | 71.0526 | 65.5172 |
|  | 0.02 | 69.2982 | 72.4138 |
| Pybel.FP4-BCS-F-0.1-1-I.II-III.IV | 0 | 61.5385 | 62.5000 |
|  | 0.001 | 63.8462 | 53.1250 |
|  | 0.005 | 70.7692 | 50.0000 |
|  | 0.01 | 73.8462 | 68.7500 |
|  | <u>0.02</u> | <u>77.6923</u> | <u>75.0000</u> |
| Pybel.FP4-BCS-F-1-I.II-III.IV | 0 | 69.3878 | 41.6670 |
|  | 0.001 | 63.2653 | 66.6667 |
|  | 0.005 | 59.1837 | 83.3330 |
|  | 0.01 | 59.1837 | 83.3333 |
|  | <u>0.02</u> | <u>71.4286</u> | <u>91.6667</u> |

**Table 16.** SVM values of I, III-II, IV comparison after classification based on FP4 type BCS dose coefficient. Abbreviations: BCS, Biopharmaceutical classification system; SVM, Support-Vector Machine.

| **Group** | **Threshold** | **Training (%)** | **Test (%)** |
| --- | --- | --- | --- |
| Pybel.FP4-BCS-F-0-0.1-I.III-II.IV | 0 | 72.8070 | 75.8621 |
|  | 0.001 | 74.5614 | 68.9655 |
|  | 0.005 | 75.4386 | 65.5172 |
|  | <u>0.01</u> | <u>73.6842</u> | <u>72.4138</u> |
|  | 0.02 | 79.8246 | 62.0690 |
| Pybel.FP4-BCS-F-0.1-1-I.III-II.IV | 0 | 73.8462 | 68.7500 |
|  | 0.001 | 75.3846 | 56.2500 |
|  | 0.005 | 76.9231 | 68.7500 |
|  | 0.01 | 76.1538 | 62.5000 |
|  | 0.02 | 74.6154 | 59.3750 |
| Pybel.FP4-BCS-F-1-I.III-II.IV | 0 | 57.1429 | 50.0000 |
|  | 0.001 | 59.1837 | 41.6667 |
|  | 0.005 | 73.4694 | 50.0000 |
|  | 0.01 | 81.6327 | 66.6667 |
|  | 0.02 | 73.4694 | 58.3330 |

### Molecular fingerprints characteristic of BCS classes III and IV

Based on the IG value, f value, dose coefficient F, and SVM prediction results, characteristic molecular fingerprints were screened for small-molecule drugs assigned to BCS classes III and IV (Tables 17-19). Feature structures of class III drugs when the dose coefficient was 0-0.1 included Secondary aliphaticamine, Tertiary_amide, Primary_alcohol, and Carbonic_acid_derivatives. A dose coefficient of 0.1-1 included Primary_alcohol and Hetero_N_basic_no_H, and a dose coefficient greater than 1 included Secondary_carbon and Michael_acceptor. Structural features of class IV drugs when the dose coefficient was 0-0.1 included Sulfonamide, Sulfonic_derivatives, and Dialkylether. A dose coefficient of 0.1-1 included NOS_methylen_ester_and_similar, Hetero_methylen_ester_and_similar, and Isothiourea. Finally, a dose coefficient greater than 1 included Sulfonamide Sulfonic_derivatives, Secondary_amides, and Vinylogous_amides.

**Table 17.** Characteristic molecular fingerprints with dose coefficients in the range of 0-0.1. Abbreviations: BCS, Biopharmaceutics Classification System; p_0_(t), the probability of fingerprint in the first category; p_1_(t), the probability of fingerprint in the second category; IG, Information gain.

| BCS category | Fingerprint | Structure name | $p_0(t)$ | $p_1(t)$ | IG |
| --- | --- | --- | --- | --- | --- |
| III | Fingerprint25 | Secondary aliphatic amine | 0.2368 | 0.0588 | 0.0386 |
|  | Fingerprint101 | Tertiary_amide | 0.1316 | 0 | 0.0514 |
|  | Fingerprint13 | Primary_alcohol | 0.1579 | 0.0588 | 0.0154 |
|  | Fingerprint143 | Carbonic_acid_derivatives | 0.1579 | 0.0588 | 0.0154 |
| IV | Fingerprint211 | Sulfonamide | 0 | 0.1176 | 0.0638 |
|  | Fingerprint214 | Sulfonic_derivatives | 0 | 0.1176 | 0.0638 |
|  | Fingerprint16 | Dialkylether | 0.1053 | 0 | 0.0406 |

**Table 18.**
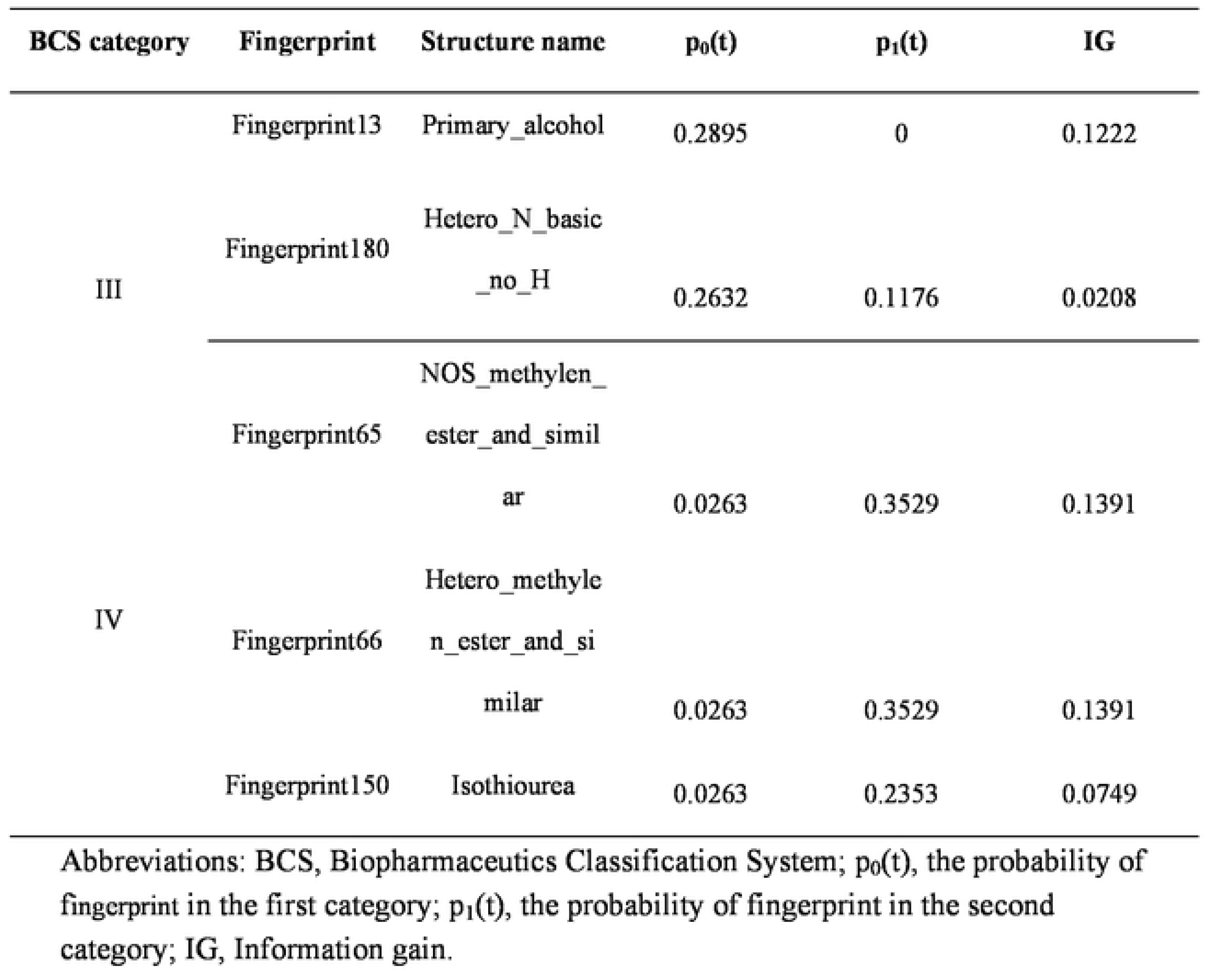
Characteristic molecular fingerprints with dose coefficients in the range of 0.1-1. Abbreviations: BCS, Biopharmaceutics Classification System; p_0_(t), the probability of fingerprint in the first category; p_1_(t), the probability of fingerprint in the second category; IG, Information gain.

**Table 19.** Characteristic molecular fingerprints with dose coefficients greater than 1. Abbreviations: BCS, Biopharmaceutics Classification System; p_0_(t), the probability of fingerprint in the first category; p_1_(t), the probability of fingerprint in the second category; IG, Information gain.

| BCS category | Fingerprint | Structure name | $p_0(t)$ | $p_1(t)$ | IG |
| --- | --- | --- | --- | --- | --- |
| III | Fingerprint2 | Secondary_carb<br>on | 0.2895 | 0.0588 | 0.0573 |
|  | Fingerprint303 | Michael_accept<br>or | 0.0789 | 0 | 0.0301 |
|  | Fingerprint211 | Sulfonamide | 0 | 0.1765 | 0.0976 |
|  | Fingerprint214 | Sulfonic_derivat<br>ives | 0 | 0.1765 | 0.0976 |
| IV | Fingerprint100 | Secondary_amid<br>es | 0.0526 | 0.2353 | 0.0483 |
|  | Fingerprint138 | Vinylogous_ami<br>des | 0.0789 | 0.2353 | 0.0313 |

### Verification of results

We verified the accuracy of our results by comparing the obtained BCS feature structures with those of small drug molecules with clear BCS classification in the BCS database (S3 File). These data confirmed that the molecular feature structures we identified were consistent with small-molecule drugs found in BCS classes III and IV, and that this method can be used as an accurate predictor.

## Discussion

In this study, we used computer simulation to analyze the common molecular fingerprints of 411 drug molecules. We determined that IG value, f value, and dose coefficient F were all necessary for the accuracy of SVM classification prediction. Lastly, we generated five macros to calculate and classify BCS (S4 File).

Different BCS classifications have been developed by the World Health Organization, the US Food and Drug Administration and the European Medicines Agency. The FDA definition of high solubility is that the highest dose allowed for a single drug administration can be dissolved in 250 ml or less of an aqueous solution at 37℃ and pH1.0-7.5. The other regulatory bodies narrow the pH range to 1.2-6.8 or 1.0-6.8 [23]. In this study, we followed the classification from the World Health Organization.

The introduction of the dose coefficient into the classification prediction software improved the reliability and accuracy of the results. Some drugs have a maximum dose of 1000 (Cefmetazole), while some drugs have a maximum dose of only 0.003 (Alfacalcidol). Moreover, small-molecule drugs have different molecular masses. In order to eliminate these two differences and make the results more reliable and accurate, we defined the concept of dose coefficient F as a standard for BCS secondary classification for the first time, which considers the ratio of the molecular mass of each small-molecule drug to the mass in the maximum dose.

This study failed to obtain the characteristic fingerprints of class I and class II drugs, which may be due to the fact that the constructed database is not large enough, such that the frequencies of fingerprints of various features are too low. Therefore, in future studies, it is necessary to continuously increase the number of small-molecule drugs in the database, so that the characteristic fingerprints obtained are more meaningful. Since the present study used only CLogP standards, future studies should include ALogP and KLogP [24] to make the research more extensive.

## Conclusion

In this study, we used FP4 fingerprints to describe the chemical structures of drugs, and calculated IG values and f values as indicators of computer simulations. Furthermore, we used SVM as the evaluation criteria for the accuracy of BCS classification. The structural features of drugs in BCS classes III and IV were successfully obtained, including Secondary aliphaticamine, Michael_acceptor, Isothiourea, and Sulfonamide Sulfonic_derivatives. These structural features can be used for the classification and formulation of drugs in these two classes. We believe that with the increase in number of the exact classification of class I and class II drugs, the characteristic structures of the two types of drugs can be obtained successfully and further guide the development of new drugs.

## Acknowledgment

This research did not receive any specific grant from funding agencies in the public, commercial, or not-for-profit sectors.

## Supporting information

**S1 Table. BCS database.**

**S1 File. IG value and the f value classification results(4 tables).**

**S2 File. Dose coefficient F classification results(18 tables).**

**S3 File. Verification results (2 tables).**

**S4 File. Calculation formulas (macro) (5 formulas).**

